# Predicting long and short-term species persistence following invasion of ecological communities

**DOI:** 10.1101/2025.04.16.649095

**Authors:** Ankit Vikrant

**Affiliations:** Department of Space, Earth and Environment, Chalmers University of Technology, Maskingränd 2, 412 58 Gothenburg, Sweden

**Keywords:** Invasions, coexistence, high-diversity

## Abstract

Large ecological communities are underpinned by a complex web of interactions whose exact structure, strength and signs are tremendously elusive. While many large-scale patterns in such communities are well-understood and predictable, the outcomes of processes such as invasions are still difficult to analyze. Given that the assembly history of communities is marked by invasion events at many points in time, it is useful to identify those aspects of invasions that can be reliably predicted even if the exact invasion outcomes cannot be determined. We introduce the notion of ‘proximate uninvadable systems’ and use these to develop a framework for predicting the structural outcomes of invasion events, i.e., what species are present/absent in the eventual equilibrium. The method is particularly illuminating in large ecological communities and applies even when the invading species is initially abundant. We test this method on a broad class of settings and also demonstrate its robustness against imperfect knowledge of species interactions. Using an example of a large food-web from peri-Alpine lakes, we show how this framework can be applied to systems with fluctuating species abundances. Given that these systems exhibit large fluctuations for prolonged periods of time, we make forecasts for extinction risk in the short term thereby extending the purview of our predictive apparatus.

## 1 Introduction

Observed ecological communities emerge from dispersal and assembly histories of species as well as the effects of the environment on them. Some assemblages are highly contingent on the assembly paths and might be impossible to build using simple bottom-up approaches — i.e. sequential introduction of species. Examples include experimental microbial communities whose constituent species pairs almost always fail to coexist (Chang et al. (2023)). Most real world examples of engineering desired communities start from existing ecosystems that are subject to species introductions — the human gut microbiome for example. This necessitates understanding community assembly through invasion outcomes.

Many theoretical studies partition invasion outcomes into broad categories such as coexistence, turnover, and invasion failure (Case (1990); Arnoldi et al. (2022). Turnover results in one or more resident species being replaced which is harder to predict than other categories of outcomes (Arnoldi et al. (2022)). This limitation also precludes identifying resident species that go extinct. Identity agnostic metrics such as species richness have dominated many areas of ecology, particularly community assembly.

While discerning the composition of an invaded community might not be possible or even sought after, metrics such as species richness belie the consequences of invasion processes. Invasions might lead to turnover and preserve the number of species, but species composition could still be altered via effects like species homogenization (Dornelas et al. (2014)). A possible risk is the loss of some functional aspects of the ecosystem, which calls for a shift toward functional diversity indices (Renault et al. (2022)). In other contexts like biodiversity conservation, species identities are even more relevant. The structure of an ecological community is possibly the simplest way to capture species identities, which further encodes assembly histories during sequential invasion events.

The goal of this paper is to predict structural outcomes of invasions, i.e., what species are present/absent in the final community following an invasion event. Our approach is based on the concept of uninvadable equilibria. The study of uninvadable equilibria is rooted in evolutionary game theory — specifically the notion of evolutionary stable strategies (ESS). Consider the replicator equations that describe the dynamics of relative frequencies *p*_*i*_ of strategies or species:

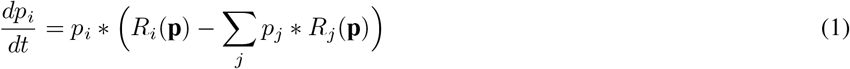

where *R*_*i*_ is the fitness for each species that depends on interactions with other species. The second term in the bracket is the mean fitness over all species. The simplest form of the fitness is *R*_*i*_ = ∑_*j*_ *A*_*ij*_*p*_*j*_ where *A*_*ij*_ is the interaction strength of a given species pair. To keep the notation short, we’ll write these terms in a vector notation such that the vector of fitnesses can be written as **R** = *A***p**. *A* is the interaction matrix.

A feasible equilibrium of the replicator equations is called a mixed strategy, which is a distribution over species frequencies. Consider the case where a resident population exists at a mixed strategy distribution **q**. If the population always reverts back to the distribution **q** whenever a small fraction *ϵ* of the residents is replaced by an invading population with a different distribution **p**, then **q** constitutes an ESS. This requires the residents to have greater fitness than all **p**, i.e.:

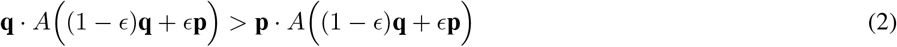

The following two conditions could therefore define an ESS:

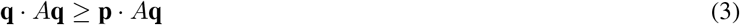

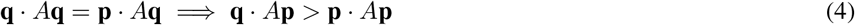

A stronger version of the ESS can be defined by replacing 4 by:

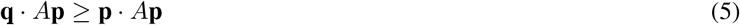

There are various ways of interpreting the ESS definitions, and we will focus on the one that results by rewriting equation 2. Note that the distribution **r** = (1 − *ϵ*)**q** + *ϵ***p** is sufficiently close to **q**. Then equation 2 follows from the following condition:

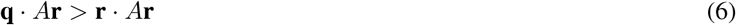

Therefore, an equilibrium **q** is strongly uninvadable if for any **r** sufficiently close to **q, q** · *A***r** *>* **r** · *A***r** holds.

## 2 Methods

Figure 1A summarizes the general prediction workflow used in this paper.

**Figure 1:**
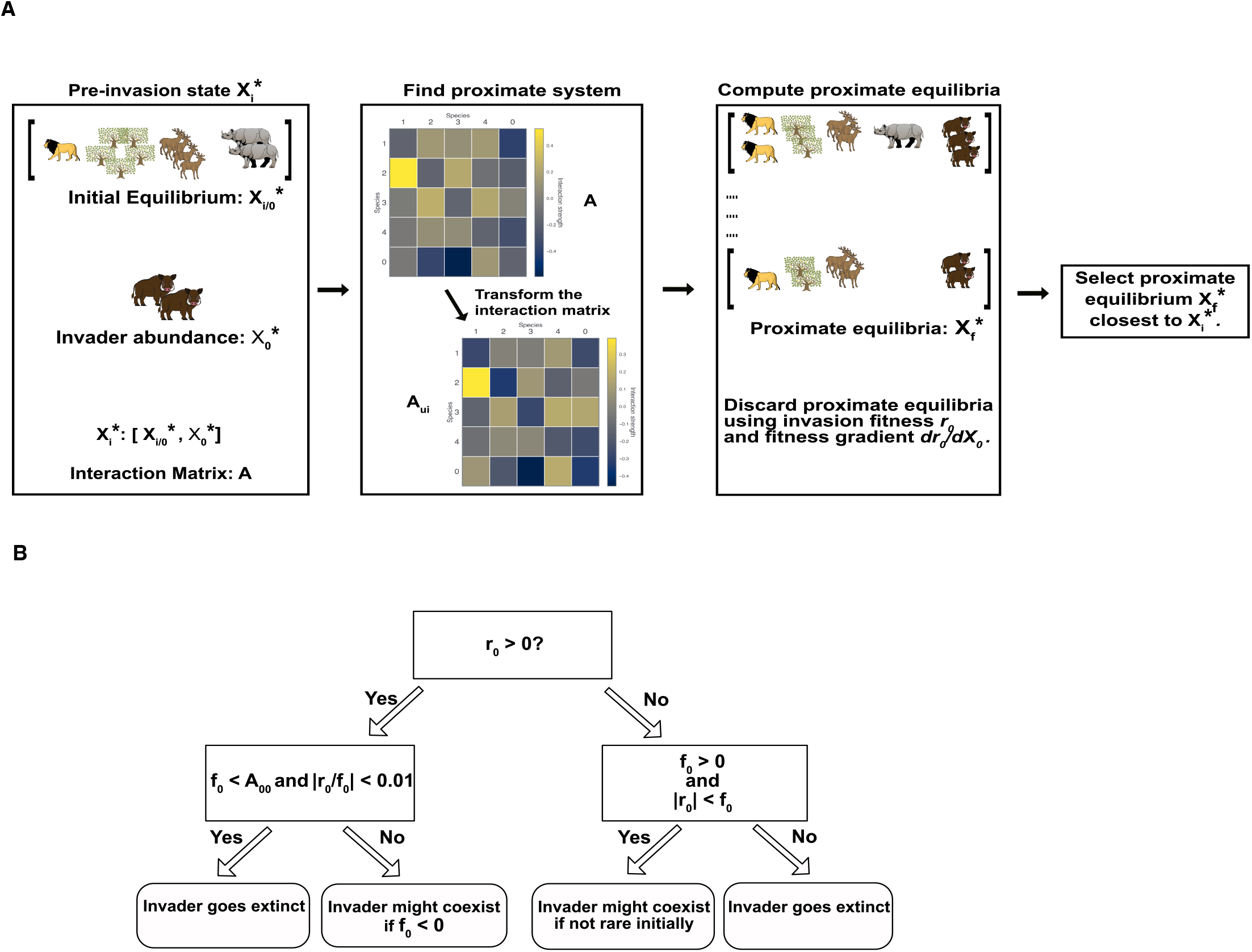
**A.** This illustration shows our prediction workflow in the most general case. The proximate equilibria here differ in terms of having different set of surviving species. As we discussed in the Methods section, a possible estimate for the invader’s final abundance can also be used to discard certain outcomes for type I functional response. **B**. The decision tree determines the outcomes for the invader based on the values of the invasion fitness *r*_0_ and the fitness gradient *f*_0_. The invader always goes extinct if its initial abundance is low and *r*_0_ *<* 0.

We consider population dynamics of the following form:

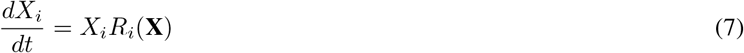

where *R*_*i*_(**X**) is the effective growth rate function that usually depends on inter-species interactions.

We only focus on stable equilibria that result from the dynamics of the resident community and the invader. There could be many possible equilibria, each characterized by a combination of species that persist. The post-invasion state is the actual equilibrium reached by the system. Our goal is to predict the structure of this post-invasion state — i.e. identify the set of species that persist at equilibrium.

The pre-invasion state is composed of a resident community at equilibrium and an invader at some initial abundance. We define the initial abundance vector as 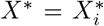 where the index *i* = 0 is denotes the invading species. Most invasion analyses consider invading species at low starting abundances such that density dependence of the invader can be ignored, but we allow for invader abundances that are not small.

### 2.1 The landscape of proximate uninvadable equilibria

The fixed points of equation 7 for which *R*_*i*_(**X**^*^) *<* 0 whenever 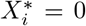 are called uninvadable equilibria or saturated rest points (Hofbauer & Sigmund (1998)). Similarly, for replicator equations the condition 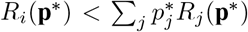 holds whenever 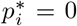. These equilibria entail that extinct species cannot reinvade if introduced in small abundances. If global stability is also endowed, then the system admits a unique uninvadable equilibrium which would also be an ESS.

Not all systems admit uninvadable equilibria, but we argue that one could use uninvadable equilibria of a nearby system to identify the eventual outcome of the invasion event. We call such nearby systems as ‘proximate systems’ and the corresponding equilibria as ‘proximate equilibria’ from hereon.

The correct proximate equilibrium would be structurally similar to the actual equilibrium in terms of the species present or absent. For GLV equations with type I functional response, if the interaction matrix *A* is such that *A* + *A*^*T*^ is negative definite, then the system is guaranteed to have an uninvadable equilibrium. Else if the largest eigenvalue of *A* + *A*^*T*^ is *λ*_*max*_ such that *λ*_*max*_ *>* 0, then the proximate system can be obtained in various ways. We use two approaches to construct such proximate systems:

1. Subtracting the following constant diagonal matrix from *A* ensures that *A* + *A*^*T*^ is negative definite :

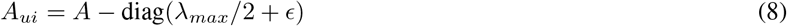

where *ϵ* is a small but positive.
2. We use a second approach based on a method to compute the nearest symmetric positive semi-definite matrix (Higham (1988)). This original method can be modified to estimate the nearest negative definite matrices as well. The implementation is discussed in Supplementary Information section S1.

### 2.2 Criteria to filter and predict invasion outcomes

The main criterion for predicting the structure of the invasion outcome is the distance between the pre-invasion state and the proximate equilibria – closer is better. We use relative entropy or the Kullback-Liebler Divergence to compute distances between normalized abundance vectors. For type I functional response, the negative definite matrix 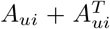 can also be used to measure distances between vectors *X*_*i*_ and *X*_*f*_ as 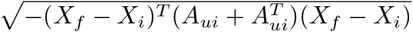.

In many cases, we can prune the space of possible outcomes before applying our main criterion. The pruning process uses the invasion fitness and its initial gradient — termed as fitness gradient from hereon. In the ecological literature, invasion fitness is called the invasion growth rate when rare, and is defined in the limit of low invader abundance. In this limit, invasion fitness is a good predictor of whether the invader survives in the final equilibrium or not.

Invasion fitness can be defined using the effective growth rate by excluding the terms that depend on the invader’s abundance, i.e.:

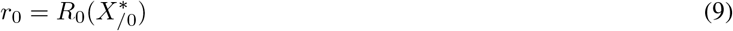

where the subscript */*0 denotes exclusion of the invader abundance. We define the fitness gradient using the formalism described in Arnoldi et al. (2022). The addition of an invading species acts as a press perturbation on the growth rates of the resident community, which results in the following expression for the fitness gradient:

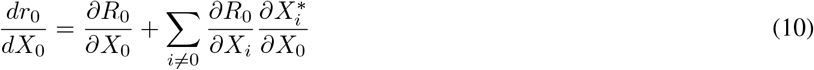

The invasion fitness and fitness gradient help infer the trend of the invader’s abundance. This abundance trend can then be used to rule out one or more proximate equilibria — for example, some values of invasion fitness and fitness gradient disallow coexistence of all species thereby ruling out a fully feasible equilibrium. We show how fitness gradient can be computed for a very general case in section S2 of the Supplementary Information.

To illustrate how outcomes are filtered, we consider communities in three different chronological states with regard to invasions: pre-invasion, transient state and post-invasion state.

We use the abundance trend of the invader to characterize the transient state. The decision tree (Figure 1B) shows certain branches along which the invader’s abundance decreases, resulting in extinction. We assume that if the invader goes extinct then the resident community goes back to its original state, and the final outcome is thus easily determined. In cases where the invader’s extinction can’t be determined, we can still filter out a large space of possible outcomes under conditions that allow for estimation of the invader’s final abundance (Figure 1B). This is possible due to a classic result from evolutionary game theory (Bomze (1991)). We describe the filtering process for the gLV system in detail in section S3 of the Supplementary Information.

We first describe the application of this framework to an evolutionary context where new mutant strains invade an existing community and the functional response is of type I. This serves as the basis of many other invasion contexts that we discuss subsequently.

### 2.3 The case of invading mutant strains

We draw pairwise interactions from a normal distribution with mean *μ/*(*c* * *N*) and standard deviation 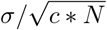 where *N* is the number of species including the invader and *c* is the sparseness of the interaction matrix. This choice ensures that the effects on any species become independent of the number of species for large *N*. We set *c* = 0.25.

We correlate the vector of invader’s pairwise interactions to that of a randomly selected resident species in the community. We set the correlation coefficient *ρ* to a high value, i.e., 0.8 for this evolutionary context. Further, the interaction from the selected resident to the invader (and vice-versa) is set to a similar magnitude as their self-interactions. More generally, for the case when the two strains have the same self-interaction term *A*_*bb*_, we can write down the interactions *A*_0*s*_ and *A*_*s*0_ as 2 * *A*_*bb*_ − *k* and *k* respectively, where *A*_0*s*_ is the effect on invader 0 from the selected strain *s*. While mathematically *k* can be any real number, it is biologically more reasonable if *k* lies sufficiently close to *A*_*bb*_. This choice ensures that if the interaction profile of the selected resident and the invading mutant were exactly the same, then the dynamics reduces to that of the original resident community. We also draw *k* from a normal distribution centred around 1.

We also allow for heterogeneity in the self-interaction term by drawing carrying capacities from a normal distribution with mean 1 and standard deviation 0.2. If the self-interactions for the selected resident and invader are *A*_*ss*_ and *A*_00_ respectively, then the interactions between two species can be set as *k* and *A*_*ss*_ + *A*_00_ − *k*.

### 2.4 The case of invading species

We draw the parameters of the invading species in two ways.

1. The first way follows directly from how the interactions are drawn for a mutant strain. We pick a random resident species to set up the interaction profile of the invader, but the correlation coefficient *ρ* is set to a lower value. We set this value to 0.5, but we also analyze cases where this correlation is much lower such that the interactions of the invader are independently drawn.
2. The second approach is the same except for how the interactions between the selected resident and the invader are drawn. We draw *A*_0*s*_ from the same distribution as the resident species. Based on this value, *A*_*s*0_ is drawn such that *A*_*s*0_ and *A*_0*s*_ admit the same correlation coefficient as that between the selected resident and the invader’s interactions. Ecologically, this approach corresponds to an invader that does not compete strongly with just one (or a few) resident species. Further, such an invader is much more novel in cases where the correlation coefficient is close to 0.

### 2.5 Invasion of top-down versus sequentially assembled communities

We want to compare the suitability of our framework to large ecological communities which are assembled in two ways:

1. The first case is that of sequentially assembled ecological communities. We start with an initial feasible community of 10 species and introduce invaders one at a time. We track the predictive ability of our framework as the starting community is pruned by subsequent invasions. We expect that such a bottom-up assembly process cannot build many larger complex communities as recent experiments have shown (Chang et al. (2023)). Therefore, we hypothesize that predicting invasion outcomes in this case should be easier than that of top-down assembled communities.
2. In the second case, we build feasible resident communities having between 10 and 100 species in a top-down manner, i.e. by introducing all species at once. For each of these communities, we introduce single invading species. For every fixed number of initial species, we generate multiple resident communities and corresponding invading species with statistically similar parameters. The results are then compared for resident communities of different sizes.

### 2.6 The effect of uncertainty in species interactions

For all cases discussed in this paper, we calculate the fitness gradient using the matrix *A*_*ui*_ from the proximate system. While this already incorporates an element of uncertainty in the systems being analyzed, we also consider cases where the interspecies interactions are imperfectly known. We again add invaders sequentially, and the predictions are made using the matrix of interspecies interactions but with some added noise. Given that our framework already allows for non-uniqueness of proximate systems, we expect our predictions to be robust to small uncertainties in the underlying interaction network. We use the method in (Higham (1988)) to find proximate systems since the off-diagonal interactions are already modified by noise, so simply subtracting constants from the diagonal might result in a proximate system that is much more distant.

### 2.7 The case of type II functional response

The consumptive interactions for type II functional response are defined using mass-based niche values. After defining an initial community of resident species, we introduced invaders by choosing random niche values to generate interactions with other species. The detailed process and the corresponding results are described in Supplementary Information section S4.

### 2.8 Evaluating predictive ability

We define the predictive score such that it takes values between 0 and 1, with 1 representing correct prediction of the invasion outcome’s structure. For cases where a fraction of the extinct species are correctly identified, the prediction score is simply the ratio of the predicted number and the actual number of extinct species. In all other cases, the prediction score is 0.

For every invasion event, we only explore proximate equilibria with up to two extinctions since the number of possible equilibria grow exponentially with the number of species which increases the computational time tremendously. Consequently, in cases when more than two species go extinct, the prediction score is always less than 1.

### 2.9 Forecasting extinction risk in an invaded food-web

We apply our framework to analyze food-web data from peri-Alpine lakes (Vagnon *et al*. (2022)). The underlying population dynamics model was parameterised using data on body sizes, metabolic rates, maximum growth rates and habitat and diet preferences of the constituent species (More details in supplementary information section S5). The population dynamics does not settle to any equilibria and shows very complicated transient dynamics with fluctuating species abundances. Many species survive at low abundances for long times, but eventually go extinct if the dynamics is run for many time steps. This motivates forecasting the effects of invading species while the dynamics is still in a fluctuating transient state. We try to predict any major structural changes such as extinctions of species driven by the invasive European catfish (*Silurus glanis*).

Since the dynamics of the food-web model does not settle down to stable equilibria, we cannot find proximate equilibria – involving subsets of species – with positive abundances. To circumvent this issue, we run the dynamics with the proximate system – or its subset – for a short time period and then use the abundances at the end times as proxies for the corresponding proximate equilibria. To simulate the invasion events, we first run the dynamics for 25000 time steps – similar to the original study (Vagnon et al. (2022)) – with the resident species only. Then we introduce the invasive species to further simulate the dynamics for 5000 time steps and forecast any extinctions. We repeat the process many times with slight variation in metabolic rates – white noise was added with variance equal to 0.1% of the metabolic rates, in line with the source study. We compare how our forecasts compare against extinctions in the respective simulations.

## 3 Results

All communities grow and undergo phases of species turnover as new invading species are introduced successively. In all cases, the results are comparable if not better if the number of species is large. All results here are for invading species with high starting abundance. For the case of top-down assembly, we also include results for low abundance.

Figure 2 shows how the predictive score changes with successive invasion events when the invader’s interaction profile is correlated to one resident species, and these two species strongly interact with each other. Under these conditions, invasion is much less successful than when such strong interactions do not exist (Supplementary Figure S2). The plots with *ρ* = 0.8 are more in line with invading mutants. Increasing the standard deviation *σ* makes the system structurally unstable. The initial resident community is reduced to a few species by subsequent invasions, and the species richness continues to stay at these low levels. We do not plot the results for such cases. We also include results for the case when the proximate system is found using the nearest negative definite matrix (Higham (1988)) in Supplementary Figure S1. When the invader’s interactions are drawn such that it does not have disproportionately strong interactions with any of the resident species — this is the second approach in the subsection ‘The case of invading species’ under Methods — we find that most invasion events result in coexistence (Supplementary Figures S2 and S3).

**Figure 2:**
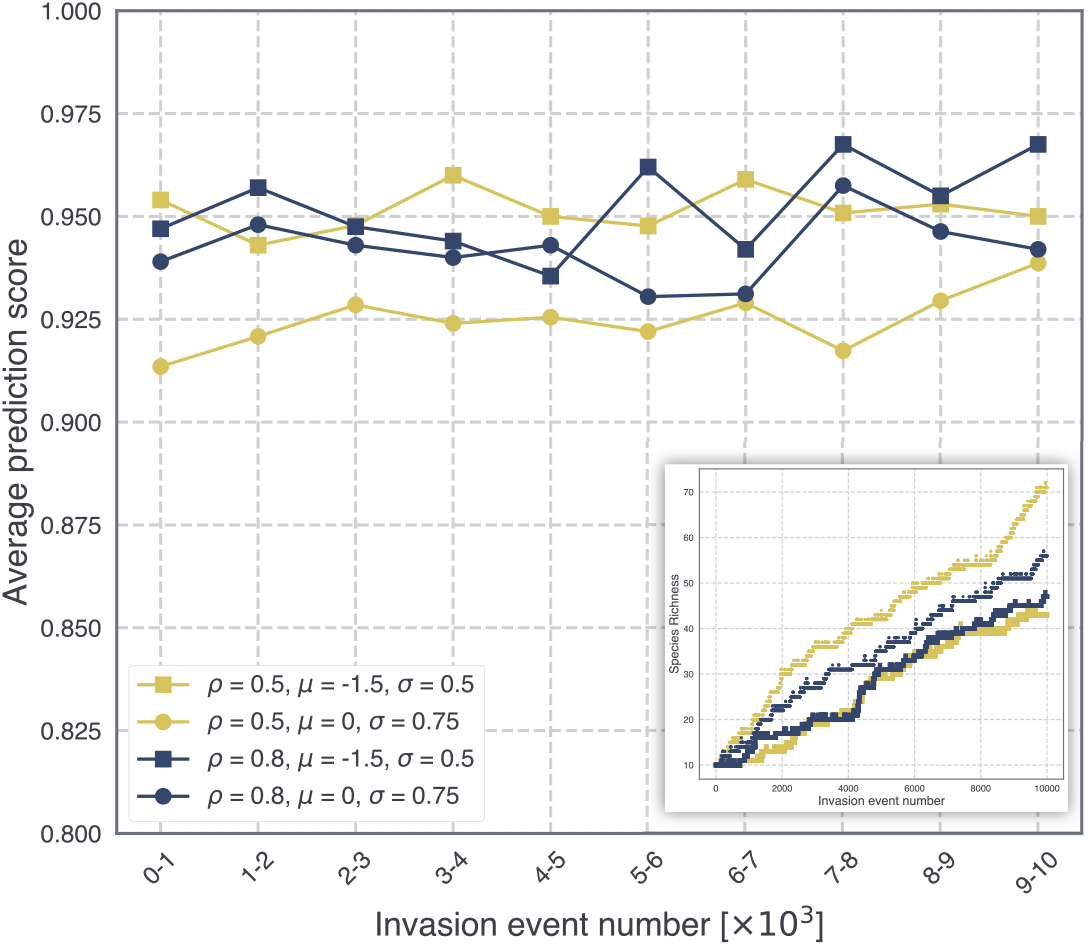
Starting with an initial feasible community of 10 species, the figure shows average prediction scores as invaders are added sequentially. The scores are averaged over 1000 consecutive invasion events. The initial resident community is generated using *μ* = 0, *σ* = 0.5 and *μ* = − 0.5, *σ* = 0.3 for the 0 and negative mean cases respectively. The inset shows the progression of species richness for each of the cases. The two lower trajectories in the inset correspond to the negative mean cases.

As expected, top-down assembly results in lower predictive scores (Figure 3). While the scores are relatively lower than sequential assembly, the framework is robust to changes in the pool size. We also simulated cases with low invader abundance, partly because this is widely studied in the literature. Here we find that prediction scores are much lower if the invader interacts strongly with one randomly selected species from the pool. In contrast, if the initially rare invader is parameterized using the second approach (check subsection ‘The case of invading species’ under Methods), the prediction scores are much higher especially for large resident communities because most events result in coexistence (Supplementary Figure S4). Since many studies analyze the case of totally random invaders with low starting abundance, we also include that case in Supplementary Figure S5.

**Figure 3:**
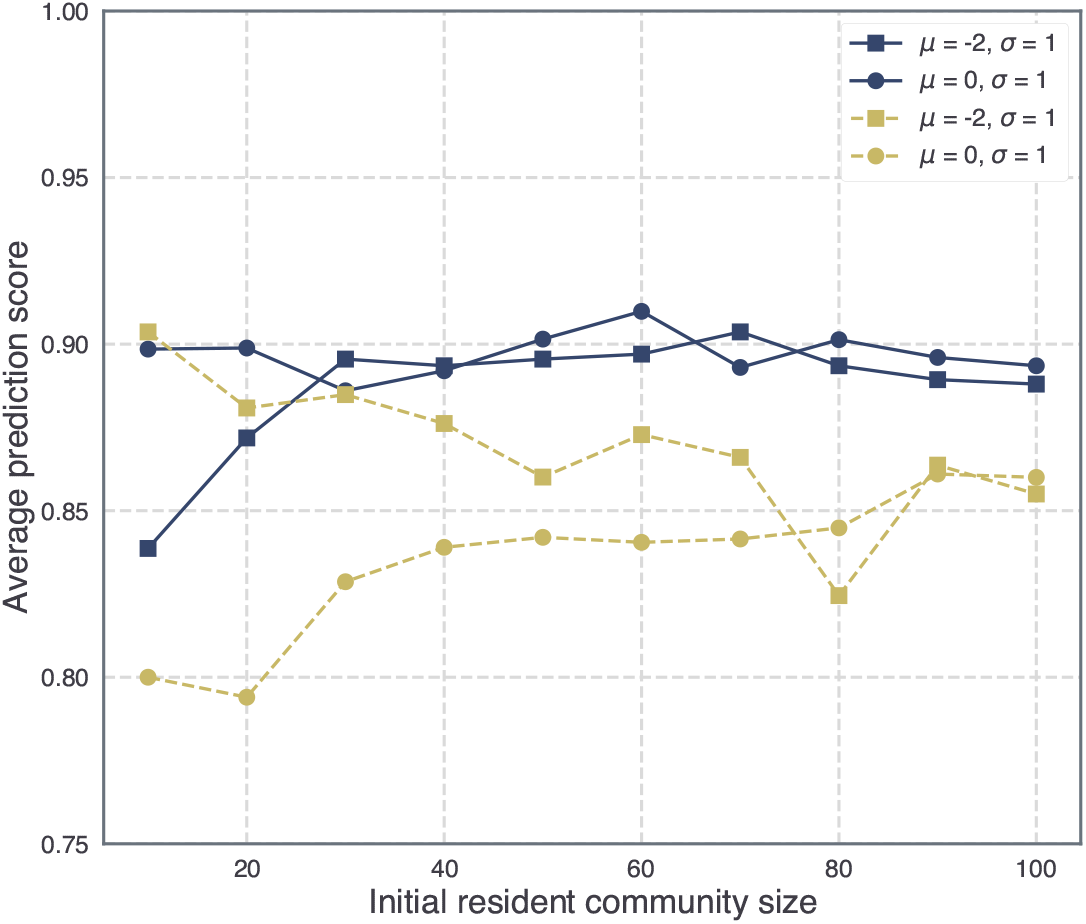
Plots for invasion of top-down assembled communities. For a given resident community size, the prediction scores are averaged over 1000 different communities with *μ* = 0, *σ* = 0.3 and *μ* = − 0.5, *σ* = 0.3 for the 0 and negative mean cases respectively. The figure legend shows the parameters of the invader’s interaction profile. *ρ* = 0.5 for all the cases shown. The dashed lines show results for initial invader abundance = 0.1.

In figure 4, we show results for cases when interactions are known imperfectly. The prediction scores are relatively lower if we add noise to all the non-zero interspecies interactions, as shown by the dashed line. In all other cases, we omit noise from the interaction terms *A*_0*i*_ which capture the effects on the invader and also appear in the invasion fitness. Under these constraints, our approach captures a large fraction of the structural outcomes except those where the invader and resident species coexist. The scores are lower for cases with mean 0 because addition of noise could change the sign of the interactions. In real contexts, we expect that the sign of these interactions could be measured reliably in many settings, and adding noise while still preserving the signs would yield better results.

**Figure 4:**
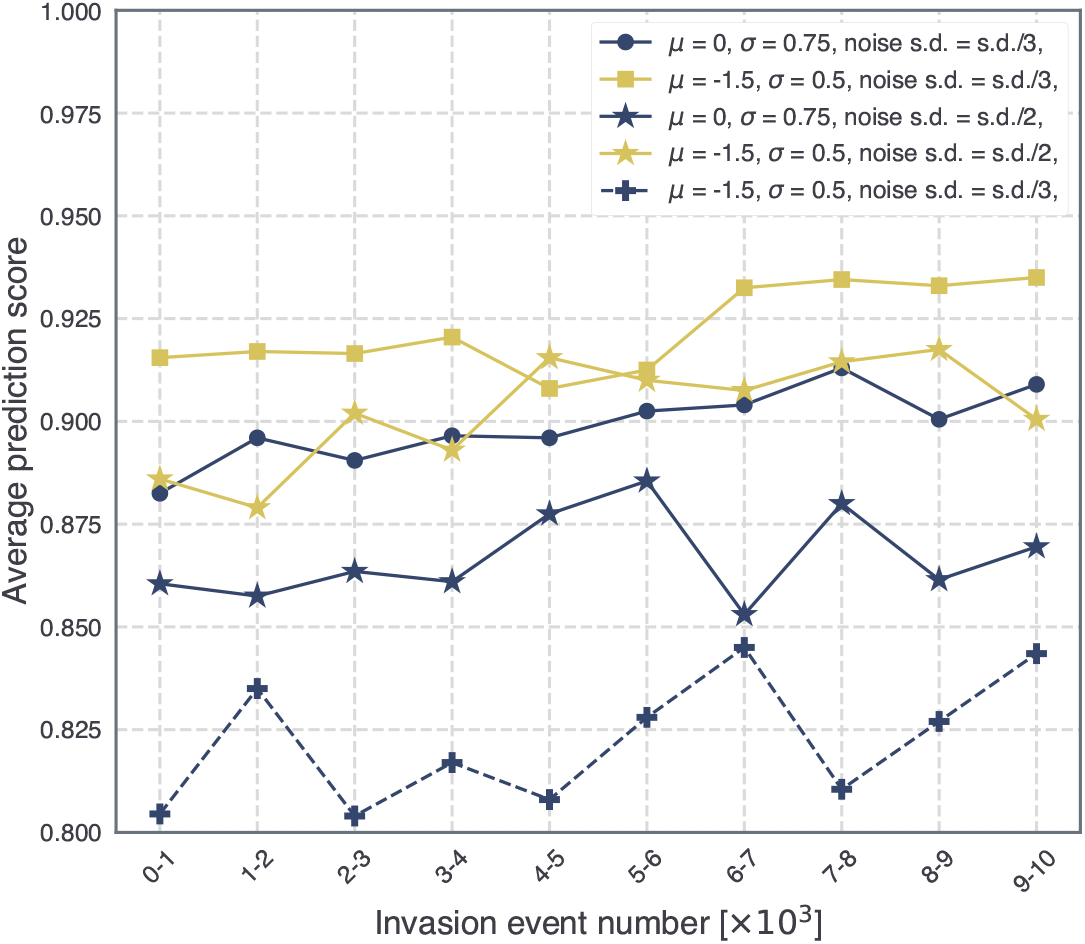
Plots for sequential assembly when noise is added to the interaction matrices. The s.d. here is equal to 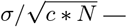 which corresponds to the distribution from which the invader’s interactions are drawn. Noise is added to the full interaction matrix only for the dashed line case. Otherwise, we assume that the effects on the invader *A*_0*i*_ are known perfectly.

We show one instance of the dynamics comparing extinct species clusters from the simulations and our forecast in Figure 5. In Figure 6, we compare the extinctions across many different iterations as the metabolic rates are slightly varied. The species cluster Daphniidae goes extinct in most realizations, which also correctly aligns with our forecast.

**Figure 5:**
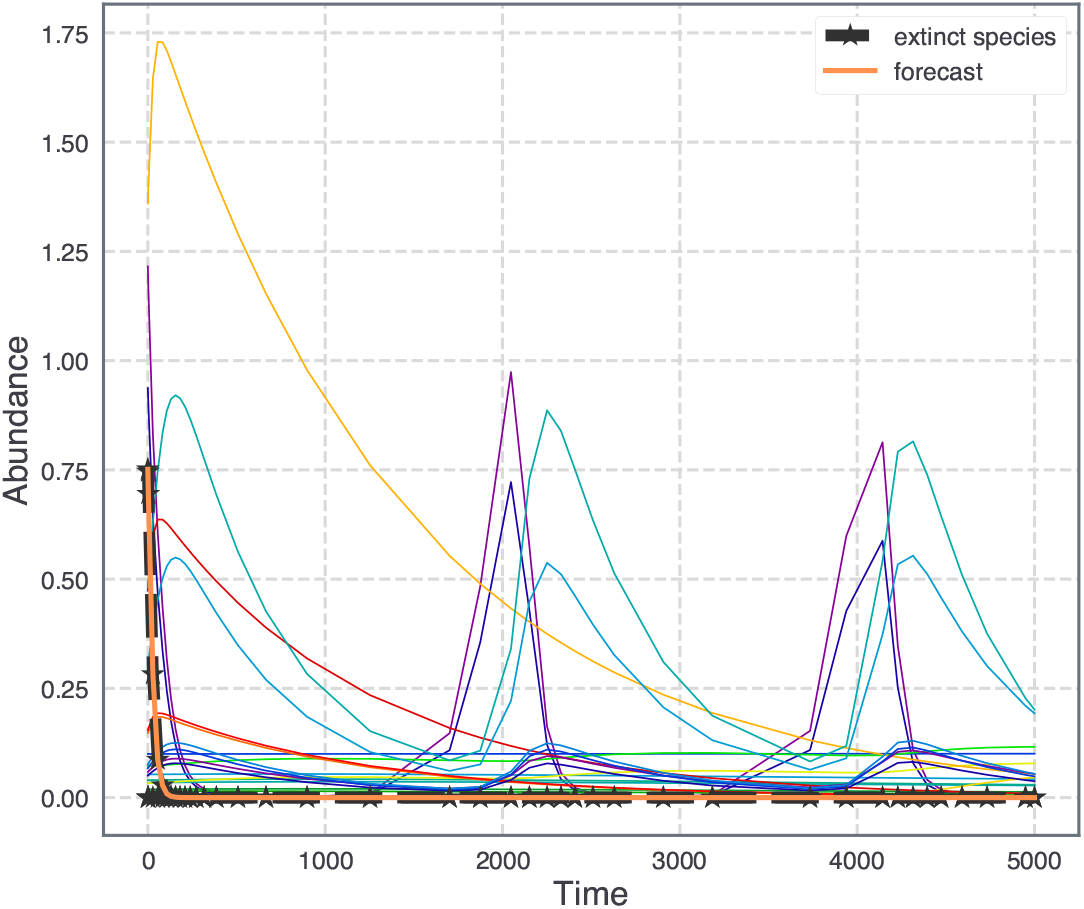
The dynamics of an aquatic food-web (Vagnon et al. (2022)) after introduction of the invasive species. As in the dynamics, our framework forecasts the extinction of the Daphniidae species cluster (thick orange line) even though its initial average abundance is high. The invasion was simulated after assembling a food-web with just the resident species.

**Figure 6:**
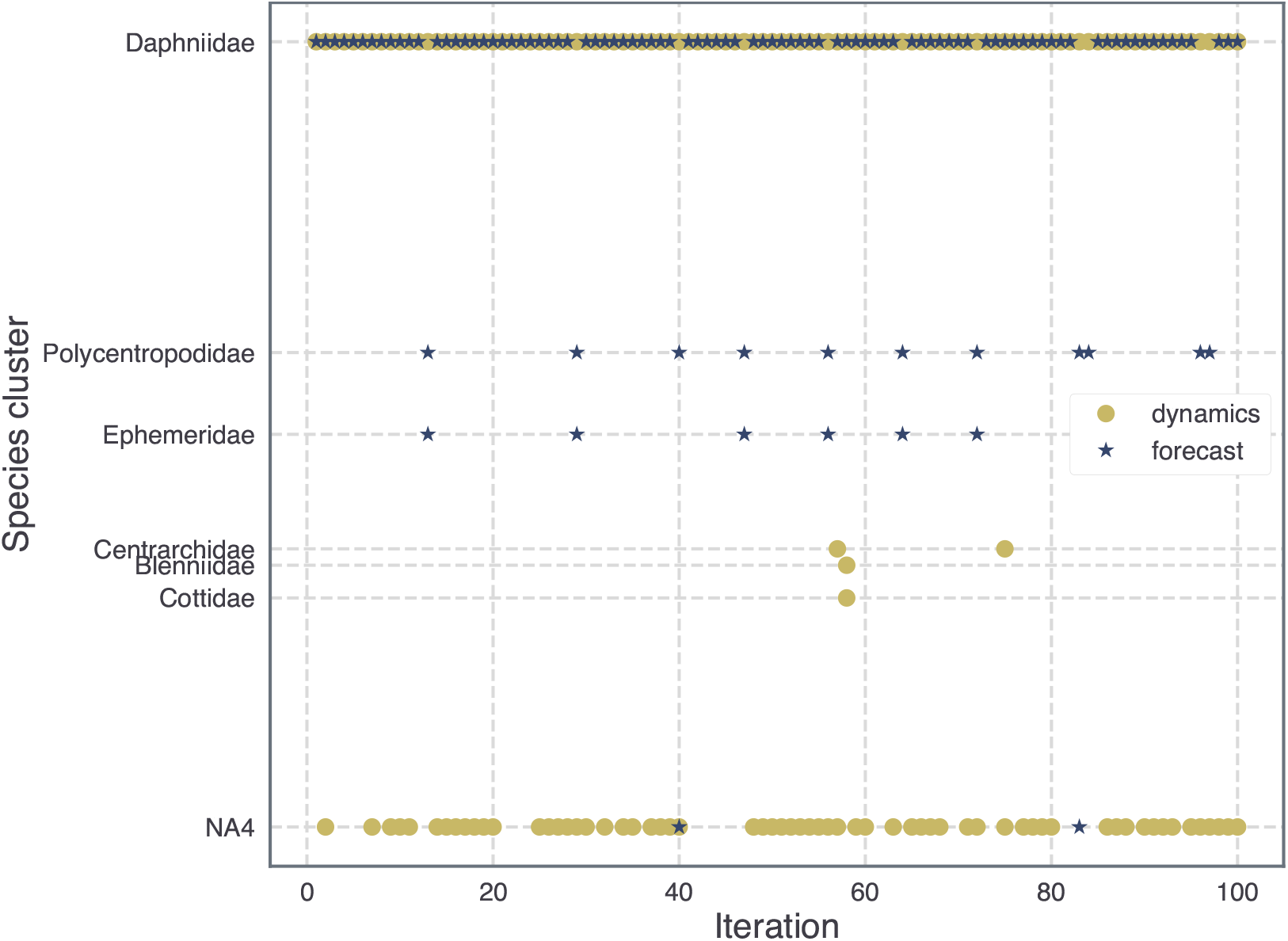
Comparisons of extinct species clusters from the simulated dynamics and our framework’s forecasts. Different iterations correspond to adding white noise to the metabolic rates of species.

## 4 Discussion

Our study proposed proximate uninvadable systems to discern the structure of invasion outcomes. We expect these proximate systems to produce better results for parameter regimes that admit greater structural stability. This is partly captured by the non-uniqueness of proximate systems, each of which correspond to different parameter values yet predict structurally similar outcomes.

We prune the space of possible outcomes by inferring the fate of the invading species. This was done by using a recent method that combines information about the invasion fitness and its gradient (Arnoldi et al. (2022)). Since our formalism is not constrained by any specific approach to discern the trajectory of the invader, one could explore other approaches to infer the same. A rough estimate of the invader’s long-term abundance can be used to evolve the system’s current state and compare its distance to the proximate equilibria, as we showed. The evolution of the system’s state could be informed by estimates of the long-term abundances of the resident species as well. We hypothesize that gauging the trend for species that experience the largest abundance changes — such as species going extinct — should be most informative in this regard.

We also analyzed the case with noisy interactions because interspecies interactions are hard to estimate reliably (Yodzis (1988); Abrams (2001); Berlow et al. (2004, 2009)). The predictive power is fairly resilient even when the interactions are known imperfectly. Interaction strengths are known to depend on species densities as well (Abrams (2001); Berlow et al. (2004)). Such dependence is captured by models with non-linear functional response such as type II — that we also tested against our method.

While our study proposes a new program for understanding the structure of equilibria, it is conceptually in line with the growing literature on structural stability in ecological systems (Rohr et al. (2014); Grilli et al. (2017)). The notion of structural stability has produced some novel insights about species coexistence and its relationship to feasibility (Grilli et al. (2017)) and predictability of invasion outcomes (Deng et al. (2024)). These results were derived using gLV systems that admit some form of global stability, such as via negative definiteness of the interaction matrix. Our work differs in terms of relaxing these constraints on the system. Also while the structuralist approach keeps the interaction matrix constant and incorporates environmental variation in the effective growth rates, the current framework identifies variations in the interaction matrix that preserve the structural outcomes.

Our results also showed that invasions are less successful if the invader interacts strongly with one or a few species (Case (1990)), and more so if the interactions are strongly correlated.

Forecasting outcomes for the invasion of peri-alpine lakes gives some key insights and early warnings. While the dynamics stays in a fluctuating state for long times, we can still forecast extinctions that are likely in the short term. The species cluster Daphniidae showed major early declines in the simulations as well as our forecasts in spite of having high average abundances in the original resident food-web. Daphniidae constitutes important zooplankton species that are one of the key food sources of resident whitefish in peri-Alpine lakes (Bourinet et al. (2023)). In a study on zooplankton communities in river Pasvik near the Barents sea, researchers documented local extinction of a large Daphnia species driven by an invasive fish species (AMUNDSEN et al. (2009)). In our analysis, we also see extinctions of most other species clusters as in Vagnon et al. (2022). There are some differences though since that paper considered top-down assembly of the food-web, whereas we first ran the dynamics with just the resident species.

Our framework has lower predictive power if invaders cause multiple extinctions in the resident community. In all of our analyses, coexistence and single extinction outcomes were more predictable than extinctions of species pairs.

We constructed the proximate systems using simple mathematical transformations that do not have a clear ecological interpretation. Given the non-uniqueness these systems, it would be useful to find transformations that optimize for some kind of ecologically meaningful proximity. Transformations that preserve the sign of interactions between species and yet produce structurally similar outcomes could be a good place to start.

## 5 Conclusion

We developed a framework that predicts structural outcomes of invasions and allows for much flexibility in initial invader abundance, structure of interactions and the functional response. Moving forward, we would like to explore the landscape of proximate systems. Possible directions include looking for metrics which place limits on how distant these proximate systems could be while still preserving the structure of outcomes.

While we did not consider space in this study, we expect that proximate systems could be constructed using heuristics such as making the self-interaction term more negative — as in the case for type II functional response. The concept of evolutionary stable strategies can be extended to other model types as well such as those with finite populations (Schaffer (1988)). Our framework could be adapted to analyse many such classes of models, and we plan to use proximate systems to investigate aspects of community assembly beyond invasions.

## Acknowledgements

I want to thank Martin Nilsson Jacobi, Susanne Pettersson and Van Savage for many useful discussions and feedback on this paper.

## Supplementary information

### S1 Finding proximate systems using the nearest negative semi-definite matrix

We first write down the eigendecomposition of *A*_*s*_ = *A* + *A*^*T*^ :

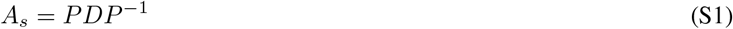

where *P* is the matrix of eigenvectors of *A*_*s*_ and *D* is a diagonal matrix with eigenvalues of *A*_*s*_ on the diagonal. We then define the matrix *A*_*snd*_ as follows:

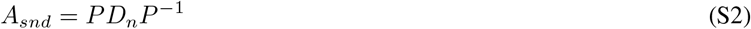

where *D*_*n*_ is constructed by replacing all the non-negative eigenvalues of *D* by small but negative values — we use −10^−15^. The proximate system is then characterized by the following matrix:

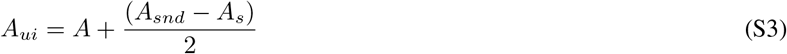

### S2 Computing fitness gradient

We can further write the indirect feedback in equation 10 as:

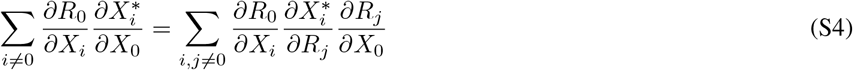

The fitness gradient can be easily computed once we can find 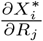. By considering the addition of an invader as a press perturbation to the growth rates of resident species, we can evaluate 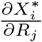 by computing the change in equilibrium abundance of resident species *i* when an infinitesimal constant term is added to the effective growth rate *R*_*j*_ of species j.

To make this explicit, consider that a species *k* gets a constant addition *I* to its growth rate. This would correspond to the system of equations:

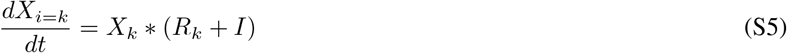

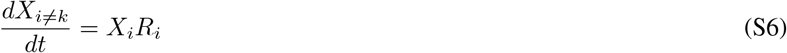

From hereon, we use the index *i* for a general resident species while *k* denotes that species whose growth rate has a constant added. The new fixed points would be obtained from 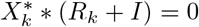 and 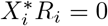. We take the partial derivatives of these equations with respect to *I*:

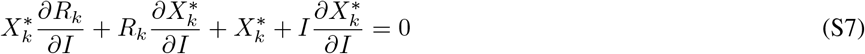

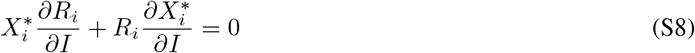

Note that for feasible equilibrium, *R*_*k*_ + *I* = 0 and *R*_*i*_ = 0 from equation S5. This reduces the set of equations S7 to

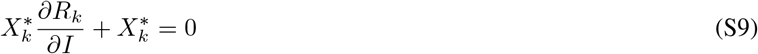

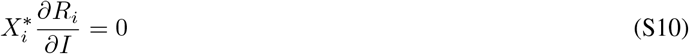

These equations can be simplified to 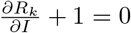 and 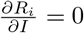. Using chain rule, we can write:

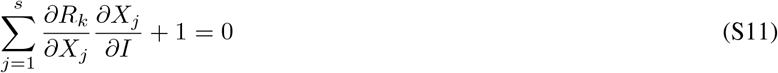

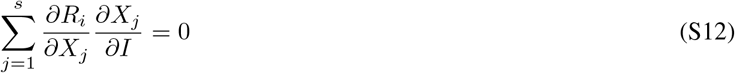

The terms 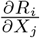 can build a matrix ***α*** whose entries are indexed by row *i* and column *j*, where the row *i* = *k* corresponds to the species for which the constant term *I* was added to the growth rate. In matrix notation, we get:

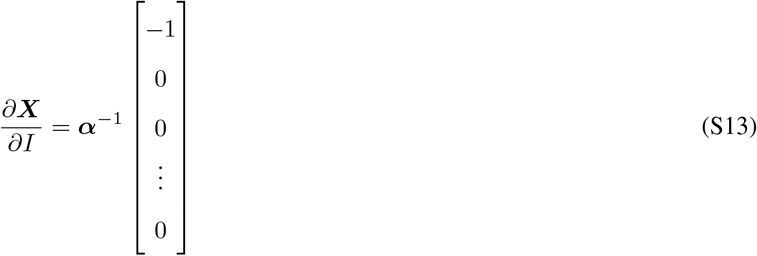

From equation S13 above, we see that the vector 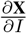 is equal to the *k*^*th*^ column of the matrix ***α***^−1^. The terms 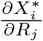 can be read off from the full matrix ***α***^−1^ to compute the fitness gradient.

### S3 Filtering outcomes using an estimate of the invader’s final abundance

Building on definition 6, it was shown that if the current state of the system is in the vicinity of a strongly uninvadable state, then the relative entropy between them decreases monotonically with time (Bomze (1991)). Relative entropy could be interpreted as a distance between two normalized abundance vectors. We want to use the result in Bomze (1991) to filter invasion outcomes by time evolving the current system state and computing its distance to the proximate equilibria.

For the gLV system, the estimation of invader’s final abundance can be divided into the following cases (Figure 2):

1. If *f*_0_ *<* 0 and *r*_0_ is not too large, the final invader abundance is approximately equal to 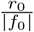 (Arnoldi et al. (2022)). We use approximation only if |*r*_0_| *<* 10 * |*f*_0_|.
2. Otherwise we set the invader’s final abundance equal to the abundance from the corresponding proximate equilibrium.

Note that the last case implies different abundance trajectories for different proximate equilibria. The trend of the abundance is inconsequential for these cases, since the distance to all proximate equilibria is guaranteed to decrease. This effectively amounts to excluding the invader abundance from the initial abundance vector and the proximate equilibria, and computing distances between these modified vectors.

In line with Bomze (1991), if the distance of a transient state to its corresponding proximate equilibrium does not decrease monotonically with time, then we filter it out.

### S4 The case of type II functional response

We consider 7 with a type II functional response such that:

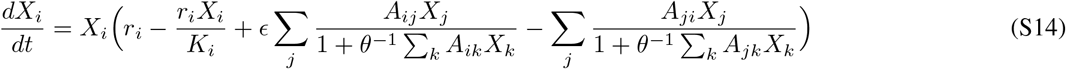

*A*_*ij*_ is the positive per-capita effect on species *i* from *j*. We draw *A*_*ij*_ based on the body masses of different species. By picking a random niche value *n*_*i*_ between 0 and 1 for each species, the body masses *w*_*i*_ are assigned as 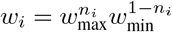 (de Roos (2021)). *As in de Roos (2021), species i* feeds on prey with niche ranging between *c*_*i*_ − *r*_*i*_*/*2 and *c*_*i*_ + *r*_*i*_*/*2, where the niche centre *c*_*i*_ is uniformly distributed between *n*_*i*_ − 2.5*/log*(*w*_*max*_*/w*_*min*_) and *n*_*i*_ − 0.5*/log*(*w*_*max*_*/w*_*min*_). The niche width *r*_*i*_ equals 1/log(*w*_*max*_*/w*_*min*_). The constants 2.5 and 0.5 ensure realistic predator-prey body mass ratios.

We set *ϵ* = 0.2 and *θ* = 1.

We could not find a simple transformation that yields proximate systems in this case, so we just make the self-interaction term more negative. The predictive power is visibly lower compared to the type I case (Supplementary Figure S6).

### S5 Food-web model of a peri-Alpine aquatic food web

We use the model in Vagnon et al. (2022) which is a population dynamics model. Many of the model parameters are allometrically scaled. The abundance of the primary producers is described by the equations:

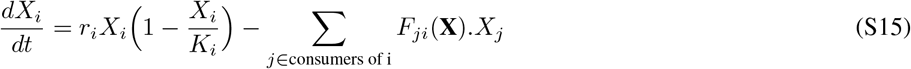

where *r*_*i*_ and *K*_*i*_ are intrinsic growth rate and carrying capacity. *F*_*ji*_ is the per capita consumption rate of consumer j on resource i, which follows a type II functional response:

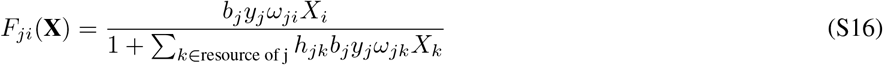

where *b*_*j*_ is the mass specific metabolic rate, *y*_*j*_ is the maximum consumption rate relative to the metabolic rate of consumer *j. ω*_*ji*_ is the proportion of resource i in the diet of consumer j, which is obtained from the allometric niche model (Vagnon et al. (2021)). h_jk_ is the handling time for resource k by consumer j. The consumer dynamics is :

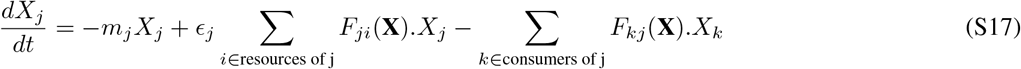

where *m*_*j*_ and *ϵ*_*j*_ are mortality rate and consumption efficiency respectively.

The relevant parameters were derived from the supplementary material of Vagnon et al. (2022), specifically data sheet 1 (R code) and data sheet 2. We used the coefficients of the feeding range stated in Vagnon et al. (2021). In allometric niche model, the feeding ranges of the consumers are found using quantile regressions between the log of predator body size and the log of prey body sizes. The lower and the upper bound are fixed at the 5% and 95% quantiles respectively. We don’t allow any trophic links between purely littoral and purely pelagic species clusters.

We add the invading catfish at low abundance of 0.1.

- **Finding the proximate system:**To find the proximate system, we subtract a small term of 0.01 times the self-interaction term for each species. While no self-regulation exists for consumers in the original model, we introduce an additional term for the proximate system only.
- **Computing proximate equilibria:** The original model stays in a fluctuating state for very long times, and if the dynamics is run even longer then most species go extinct. Instead of predicting the final outcome, we focus here on making forecasts for extinctions in the short term. Using the proximate system, we run the dynamics for 1000 time steps with one or more species removed to find proximate quasi-equilibria.

## Supplementary figures

**Figure S1.**
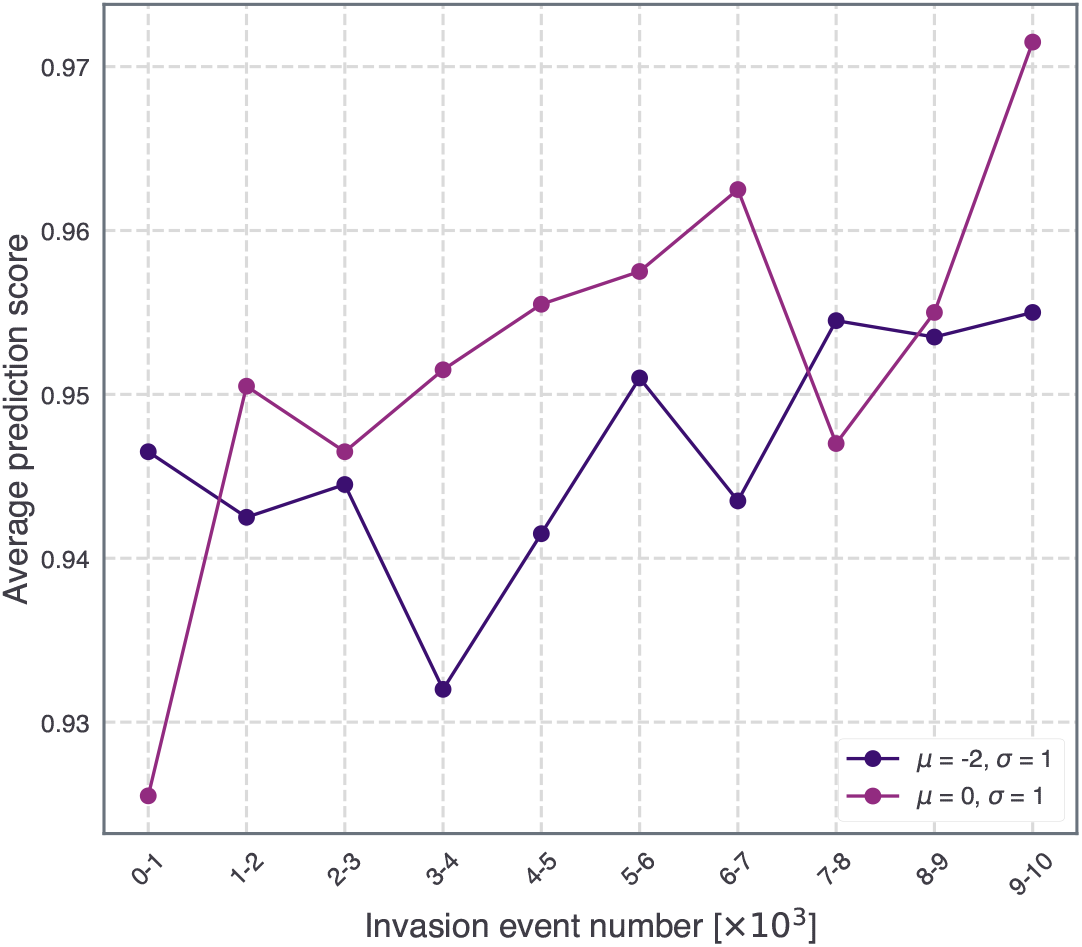
The initial resident community is generated using *μ* = 0, *σ* = 0.5 and *μ* = − 0.5, *σ* = 0.3 for the 0 and negative mean cases respectively. The proximate system is found using the nearest negative definite matrix. *ρ* = 0.5 for both cases.

**Figure S2.**
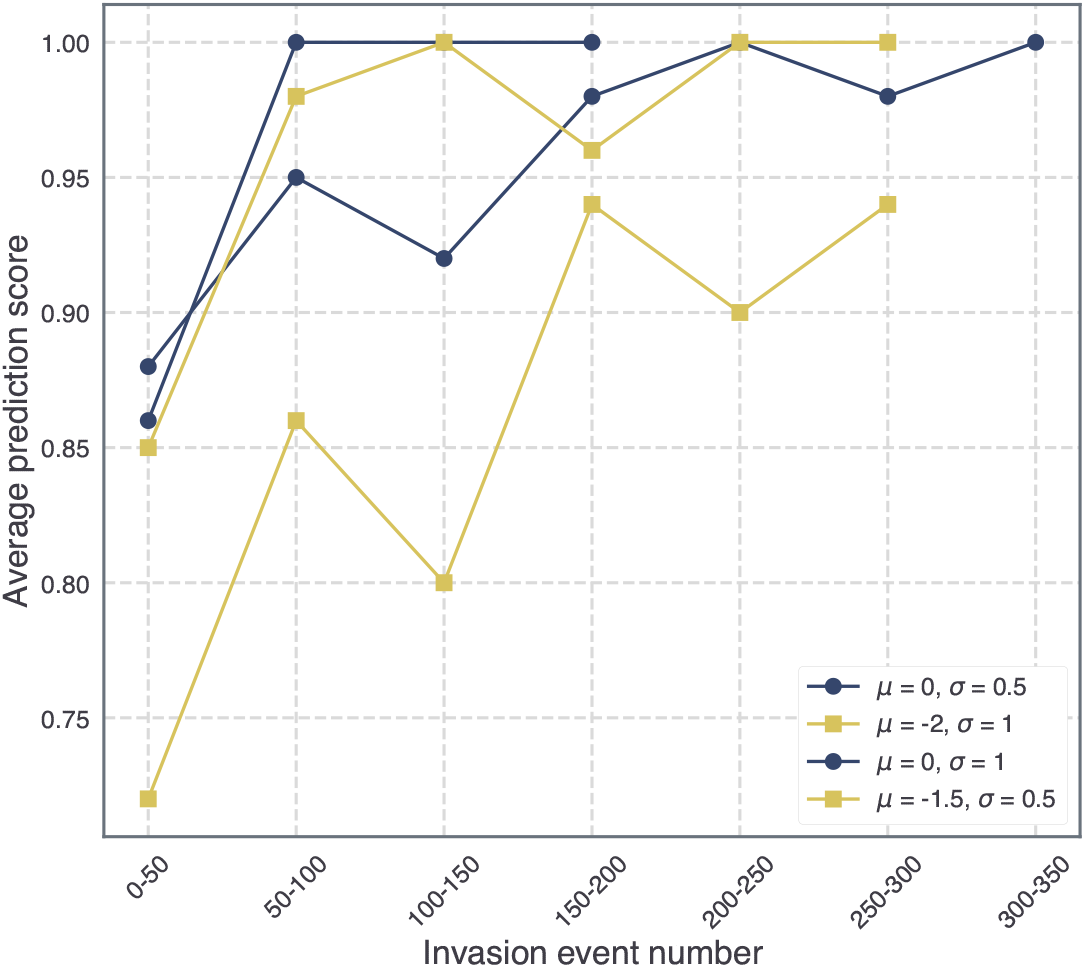
Plots for sequential invasion events when the invader’s interaction profile is correlated to a randomly selected resident species, but no disproportionately strong interactions exist. All trajectories have *ρ* = 0.5. Most events result in coexistence, and hence no more species are added after a few hundred invasion events.

**Figure S3.**
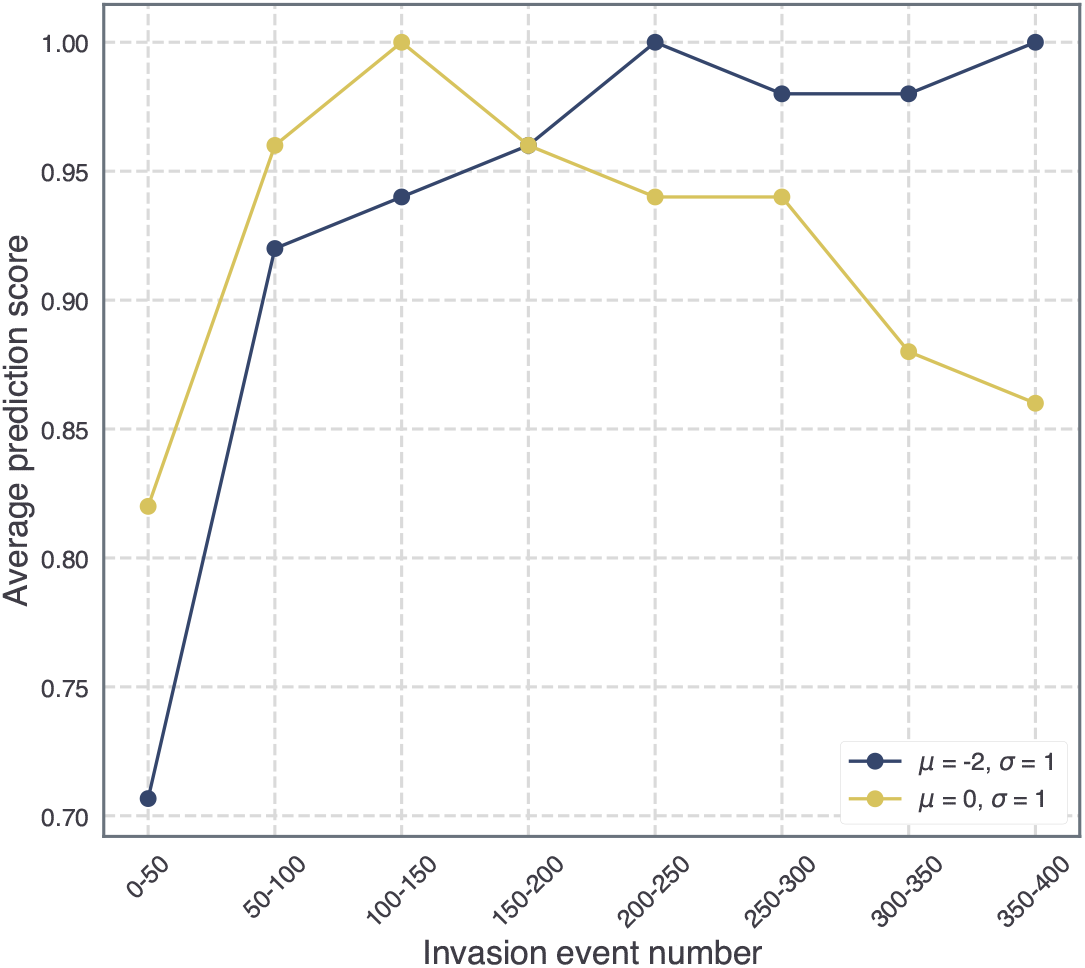
Plots for sequential invasion events when the invader’s interaction profile is totally uncorrelated from any of the resident species, i.e., *ρ* = 0. Most events result in coexistence, and hence no more species are added after a few hundred invasion events.

**Figure S4.**
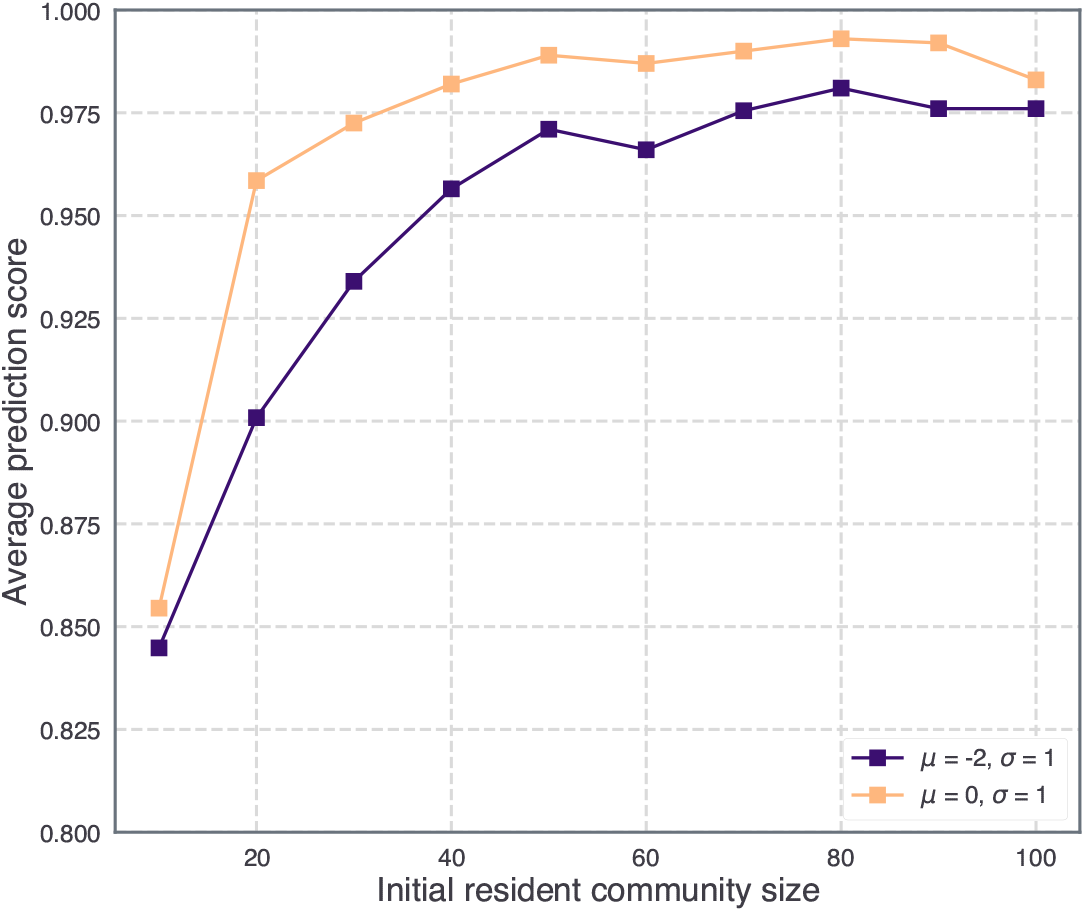
Plots for invasion of top-down assembled communities when the invader’s interaction profile is correlated to a randomly selected resident species, but no disproportionately strong interactions exist. The initial abundance of the invader is equal to 0.1 and *ρ* = 0.5. For a given resident community size, the prediction scores are averaged over 1000 initial resident communities with *μ* = 0, *σ* = 0.3 and *μ* = − 0.5, *σ* = 0.3 for the 0 and negative mean cases respectively.

**Figure S5.**
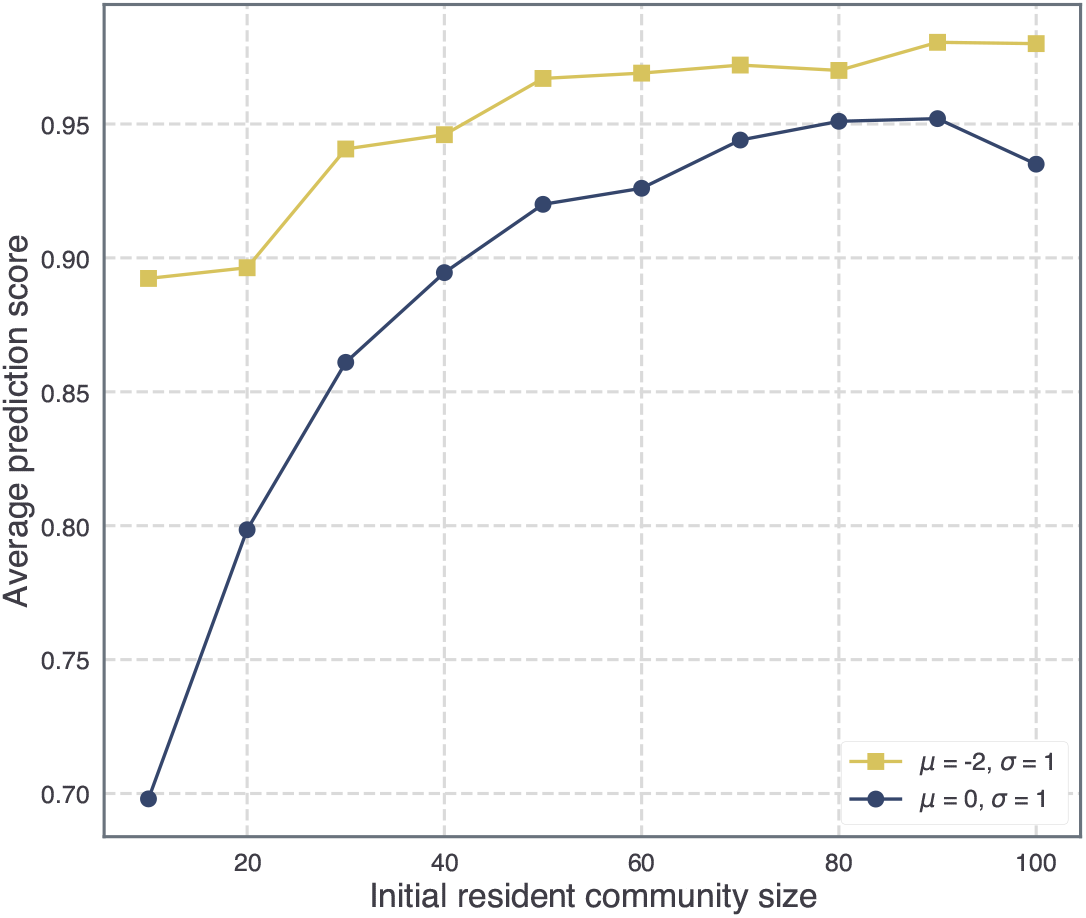
Plots for invasion of top-down assembled communities when the invader’s interaction profile is completely random (*ρ* = 0). The initial abundance of the invader is equal to 0.1. For a given resident community size, the prediction scores are averaged over 1000 initial resident communities with *μ* = 0, *σ* = 0.3 and *μ* = − 0.5, *σ* = 0.3 for the 0 and negative mean cases respectively. The negative mean case results in various outcomes involving extinctions and coexistence for up to 50 species, beyond which coexistence is more prevalent. The 0 mean case is dominated by coexistence except for the smallest pool size of 10 species.

**Figure S6.**
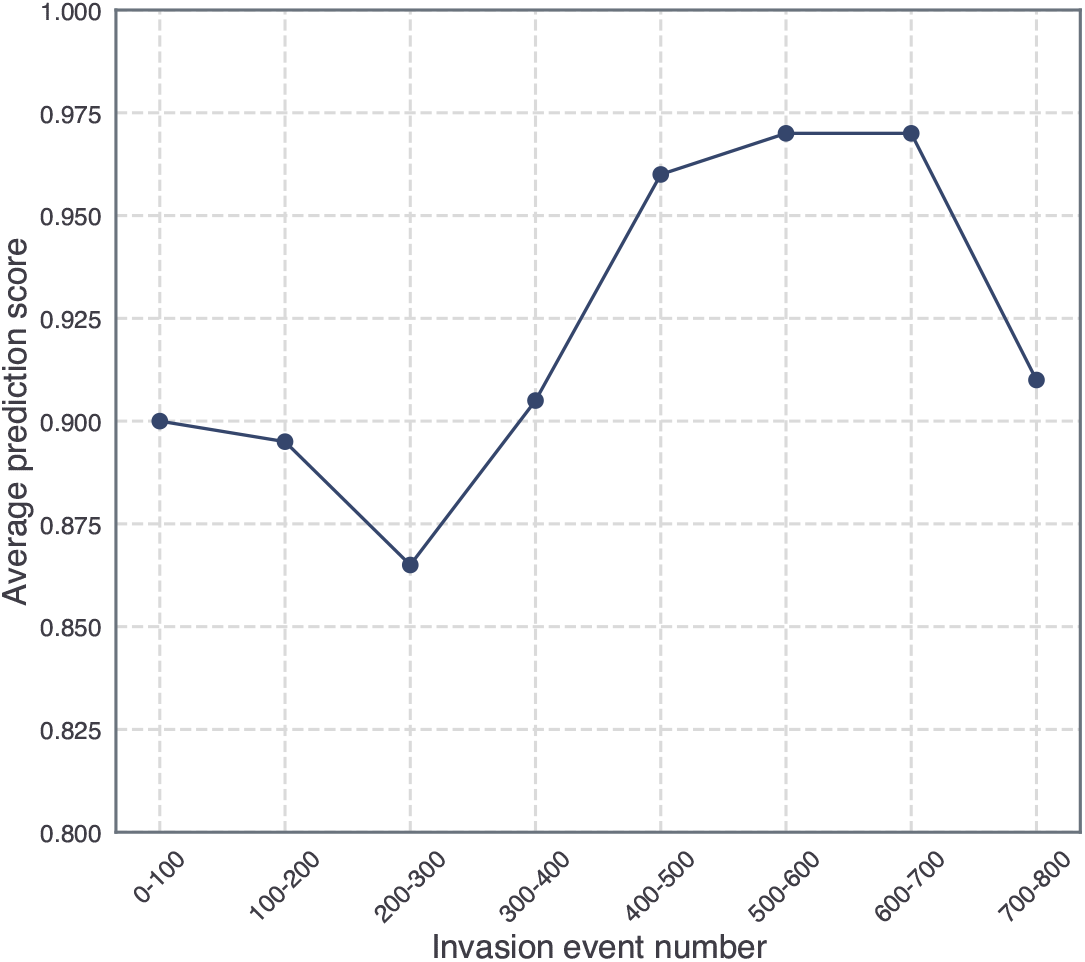
Plot for the case of type II functional response. The invader’s interactions are drawn using a random niche value as described in the Supplementary Information.

